# Emergent Programmable Behavior and Chaos in Dynamically Driven Active Filaments

**DOI:** 10.1101/2022.06.05.494577

**Authors:** Deepak Krishnamurthy, Manu Prakash

## Abstract

How the behavior of single cells emerges from their constituent sub-cellular biochemical and physical parts is an outstanding challenge at the intersection of biology and physics. A remarkable example of single-cell behavior is seen in the ciliate *Lacrymaria olor*, which hunts by striking its prey via rapid movements and protrusions of a slender neck, many times the size of the original cell body. This dynamics of the cell neck is powered by active injection of energy into this slender filamentous structure via a coat of cilia across its length and specialized cilia at the tip. How a cell can program this dynamical active filament to produce desirable behaviors like search and homing to a target remains unknown. By constructing a coupled active-elastic and hydrodynamic model of a slender filament with activity at the tip, here we uncover how cell behavior (filament shape dynamics) can be controlled via activity dynamics. Our model captures two key features of this system - dynamic activity patterns (extension and compression cycles) and active stresses that are uniquely aligned with the filament geometry - leading to a so-called “follower force” constraint. We show that active filaments under deterministic, time-varying follower forces display rich behaviors including periodic and aperiodic shape dynamics over long times. We further show that aperiodic dynamics occur due to a transition to chaos in regions of a biologically accessible parameter space. By further dissecting the non-linear dynamics of this active filament system, we discover a simple iterative map of filament shape that predicts long-term behavior. Lastly, using these iterative maps as a design tool, we demonstrate examples of how a cell could “program” filament behaviors by using frequency and amplitude modulated activity patterns. Overall, our results serve as a framework to mechanistically understand behavior in single cells such as *L. olor* and present a novel chaotic dynamical system in active elastohydrodynamics. Our work also offers a direct framework for designing programmable active matter systems using filament geometries.

**Significance statement:** Single-celled protozoa display remarkable animal-like behaviors without the aid of neurons. Mechanistically understanding how this dynamic behavior emerges from underlying physical and biochemical components is an outstanding challenge. In this work, using an active filament model, we uncover the fundamental non-linear dynamics and non-variational mechanics that underlie the complex behaviors of single cells like *Lacrymaria olor*. In doing so we discover a novel route to chaos in active elastohydrodynamic systems and the first-ever description of how chaos can drive single-cell behaviors. Lastly, we present a framework for how filament behaviors can be “programmed” using dynamic, modulated activity patterns. Overall our work provides mechanistic insights into single-cell behaviors and offers a new framework for the design of filamentous active matter systems to achieve diverse functions.

## 1 Introduction

The behavior of biological systems emerges due to the interplay of parts across molecular, cellular and tissue scales [1, 2]. Yet, despite the notion of behavior as a complex phenomena, recent efforts that quantify behavior in diverse systems have found a common theme of underlying simplicity: even seemingly complex behaviors can be described fruitfully using the dynamics in a relatively low-dimensional space, as compared to the total possible degrees-of-freedom of an organism [3, 4]. However mapping the emergence and dynamics of these finite “behavioral modes” to physical, biochemical, genetic or epigenetic features of the organism has remained elusive, primarily due to the complex intervening layers involved [5]. Unicellular ciliates, protozoans with a large number of cilia, display remarkable, rapid, animal-like behaviors such as swimming, walking, jumping and hunting without the aid of a neuro-muscular system [6, 7, 8, 9]. Thus studying ciliates presents a unique opportunity to understand emergence of complex behavior in a system with no neurons. In this reductionist approach, multiple layers of complexity of a neuro-muscular system are replaced by the relative “simplicity” of mechanical and biochemical interactions between constituent sub-cellular parts [10].

One of the most striking examples of complex behavior in a unicellular protist is seen in the predatory behavior of the ciliate *Lacrymaria olor* which utilizes a highly dynamic, ciliated, slender protrusion, that can stretch as large as ∼ 10 − 20 times the cell body size which is only ≈ 20 *µ*m. This dynamic behavior, which plays out in seconds to minutes, allows rapid and exhaustive sampling of the cell’s surroundings and enables hunting for motile prey (Fig. 1A) [11, 7]. To do so the protrusion adopts a striking range of shapes in rapid succession allowing the tip of the protrusion to visit a large fraction of the space surrounding the cell even while the cell body remains attached to a substrate (Fig. 1A) [7]. On the other hand, ciliates such as *Dileptus anser* use periodic sweeping motions of their neck-like protrusion to scan their environment [12], in a movement reminiscent of an elephant waving its trunk. These are just two of thousands of examples of single cell organisms that utilize distributed ciliary forces acting on the cell body and protrusions to achieve a diverse range of real-time behaviors [13]. When compared to better-studied cellular protrusions like filopodia which typically change length at rates of ∼ 10 *µm/min* [14], protrusions in ciliates are much more dynamic, characterized by length and shape changes occurring over millisecond and sub-second time scales [7]. The speed and reactive dynamics of these movements rule out any genetic mechanisms and pave the road to search for mechano-chemical feedback loops responsible for the diversity of behaviors.

**Fig. 1:**
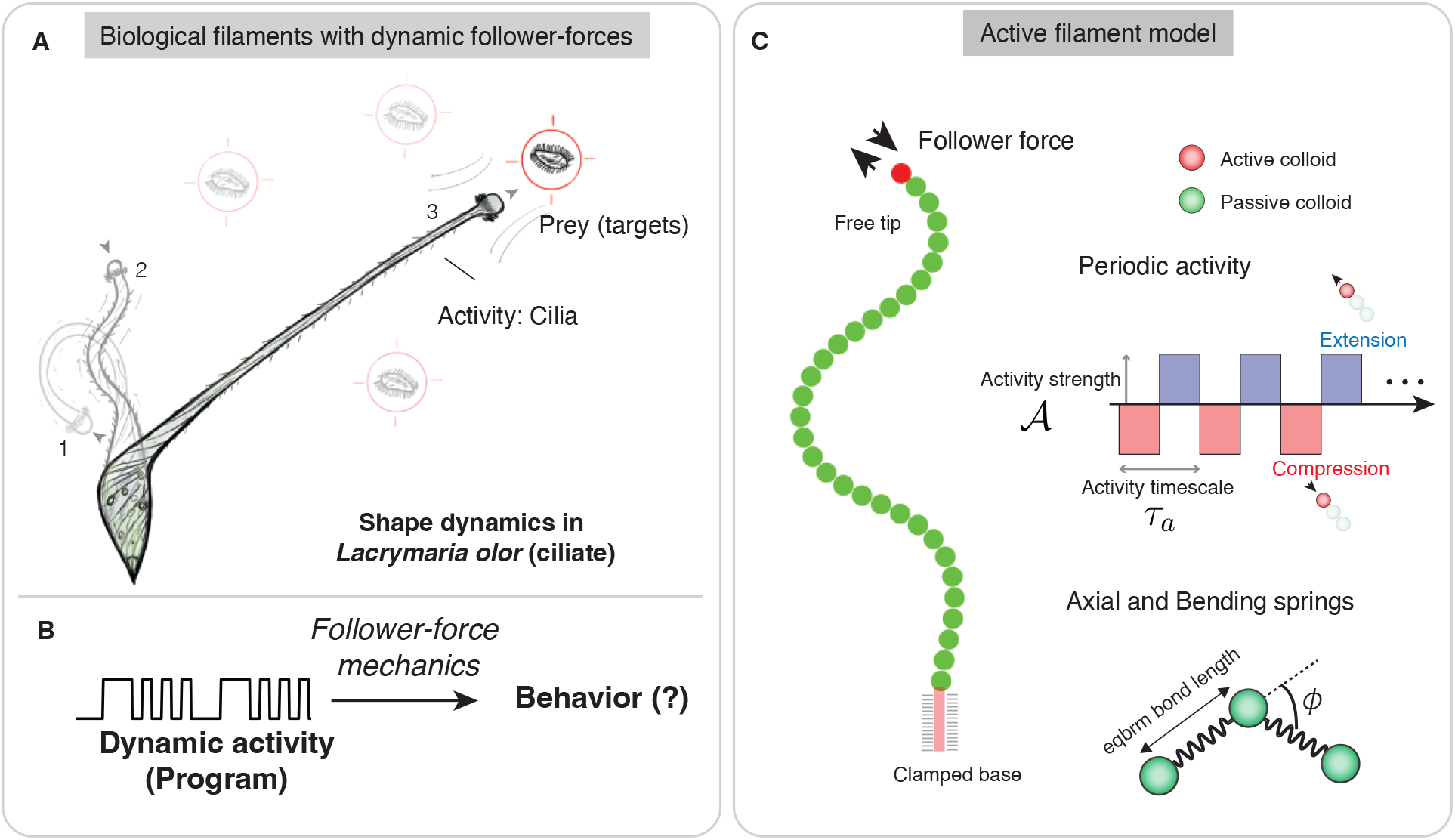
Active elastohydrodynamics of filaments under dynamic follower-forces. **(A)** The phenomena of rapid morphodynamics to search space and hunt for prey in the single-celled ciliate *Lacrymaria olor* using a long cellular protrusion. Cilia along the protrusion and specialized tip cilia provide active stresses which are constrained by the cell’s membrane and cytoskeleton and hence follow the geometry of the cell, thus giving rise to a so-called “follower-force” system. Changes in ciliary beat direction alternate the force between compression and extension. **(B)** We explore the fundamental question of how such time dynamic activity leads to observed filament behaviors due to the underlying follower-force mechanics. **(C)** Active filament model to study filament behavior under dynamic active follower-force stresses. Model filaments consist of a string of colloids which can be either active (i.e. can apply active stresses on the surrounding fluid; shown in red) or passive (green). In our work, we consider the simplest case where only the distal colloid is active (red). The activity profile is deterministic and periodic and can be parametrized by the activity strength 𝒜 and time scale for one compression + extension cycle *τ*_*a*_. Elastic potentials for inter-colloid distance and equilibrium angle constrain the filament shape along with short-ranged repulsive potentials to prevent overlap of colloids and filament self-intersection.

A key feature of protrusion dynamics exhibited by *Lacrymaria olor* is that the active stresses are constrained by the local geometry of the protrusion/neck since the cilia are directly anchored to the cell membrane and cytoskeleton. Thus the direction of forces generated by these anchored cilia change dramatically with the orientation vector of the cell surface - a classical condition in mechanics often described as a so-called “follower-force” constraint [15, 16, 17]. First described to characterize the stability of rocket engines [18], follower-forces create a unique non-conservative condition wherein the force “follows” the geometry of the system, and occur naturally in the biological context of self-driven filaments [15, 17, 16, 19]. In this work, we explore how this follower-force constraint of the filament geometry of *L. olor* encodes complex behavior leading to successful predatory characteristics of this ciliate. We further explore the role of activity dynamics to see how simply switching the direction of ciliary beat (ciliary reversal) can encode a variety of complex movements with unique organismic function.

Our recent work studying *L. olor* experimentally in lab conditions has identified key components that produce active stresses in the system - including cilia in the neck and tip of the protrusion and contractile centrin-like proteins [7]. Furthermore, the unique cytoskeletal structure of the neck imparts non-linear mechanical properties to the neck [11, 7]([20], unpublished). Although the source of active stresses and mechanical constraints has been mapped in this system, currently, we still lack a first principles understanding of how this complex behavior emerges. Specifically, how do time-varying activity patterns due to ciliary reversals, follower-force constraints and the mechanical properties of the cell lead to the observed complex shape dynamics and behavioral repertoires? What class of dynamical systems does this particular biological system fall into? Here we address these issues by leveraging the single cell context of this behavior and the relative simplicity of the filament geometry to seek an explicit deterministic framework for behavior consisting of an input, map and output, with no hidden layers: dynamic activity patterns (inputs) acting via non-conservative follower-force mechanics (map) lead to the observed time-varying shape dynamics of the cell (behavioral output) - all integrated together with functional benefits to the organism at large (Fig. 1B).

Towards this goal, in this work we explore the dynamical state space of an active filament subject to time-varying follower-forces as a toy model system to understand complex behavior of *L. olor*. Since the fundamental goal of this work is to build a modelling framework that captures the essence of the behavioral complexity observed in cells like *L. olor*, we consider the simplest model with two key features: (1) *Time-varying activity patterns*, wherein the active stresses generated by the filament can periodically change in magnitude or direction (specifically between compression and extension of the filament). This directly captures the effects of ciliary reversals which are very common across ciliates, and have previously been studied in relation to avoidance reactions in Paramecium [21]; (2) *Follower-force activity* : we constrain the active stresses to be along the local tangent vector of the filament to capture the “follower-force” constraint that arises due to the constraints on the ciliary beat plane, relative to the cell membrane and cytoskeleton (Fig. 1C). While earlier active filament models have studied the dynamics due to constant follower-force activity to try to explain the origins of, or, mimic ciliary beating, the effects of time-varying active stresses (e.g., alternating compression and extension) and how that shapes the behavioral landscape of filaments remains unknown [17]. Our goal in this work is therefore to bridge this knowledge gap and map out the space of filament behaviors under time-varying follower-forces and utilize this framework to build the first reduced-order, mechanistic model of the origins of complex behavior in a unicellular protist.

In the rest of this work we briefly describe the modelling approach and then report a rich space of behaviors in filaments under deterministic, time-varying follower-forces, including period-doubling and aperiodic filament dynamics. We show that aperiodic behaviors are due to underlying chaos in the filament dynamics, thus discovering a completely new route to chaos in viscous active elastica. To further understand and control filament dynamics we identified the fundamental non-linear features of the follower-force filament system by defining a simple iterated map based on filament shape that can predict long-term filament behaviors, including the onset of chaos. Lastly, by leveraging our insights into the non-linear filament dynamics, we also demonstrate a number of examples of “programmability” of filament behaviors by tuning biologically relevant parameters. Our work, though inspired by *L. olor*, is also very broadly applicable to the design of active matter/filament systems including microscale robots, wherein a physical task/computation, an algorithmic representation of the task, and its actual physical implementation can all be analyzed and explored systematically - with a direct analogy to Marr’s level of analysis (developed in the context of information processing systems) applied to single-celled protist behavior [22].

## 2 Results

### 2.1 Active filament model and simulation method

We consider an active filament framework [16, 23], wherein the filament consists of a series of *N* connected colloids of radius *a* that can be either active or passive (Fig. 1C). The active colloids can exert a surface slip velocity or stress on the ambient fluid [24]. The material response of the filament is captured using potentials that penalize departures from equilibrium bond length, bond angles and prevent self-intersection of the colloids. This potential is given by:

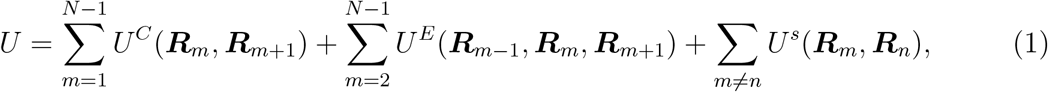

where ***R***_*m*_ is the position of the *m*^*th*^ colloid, *U*^*C*^(***R, R*′**) = *k*(*r* −*b*_0_)^2^*/*2, with *r* = |***R***−***R*′**| and ***R, R*′** correspond to the positions of two distinct colloids, *b*_0_ the equilibrium bond length, *k* the axial spring stiffness, and *U*^*E*^ = *κ*(1 −cos *ϕ*)*/b*_0_, with *κ* the bending stiffness, *ϕ* the angle between consecutive colloidal bonds (Fig. 1C). Near-field spherical and sphero-cylindrical Lennard-Jones type repulsive interactions are specified by *U*^*S*^ and prevent overlap of the colloids and also prevent self-intersection of the filament [23]. The resulting constraint force on the *n*^*th*^ colloid is given by 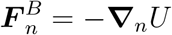.

We consider the inertialess (force and torque-free), and non-Brownian limit of the colloids corresponding to the scale of single cells (*>* 10 *µm*). The active colloid framework used captures both the local as well as many-body hydrodynamic interactions due to both passive and active surface stresses on the colloids, and results in the following equations of motion of the colloids [24, 25]:

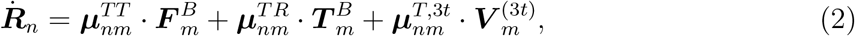

where the dot signifies temporal derivatives, 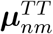 and 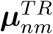 are the many-body forms of the well-known translational and rotational mobility tensors for spheres in an unbounded fluid [26], and ***µ***^*T*,3*t*^ is the propulsion tensor corresponding to the active slip on the colloids, where 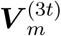 specifies the active slip due to a potential dipole of strength *d*_0_ [16, 27]. The angular degrees of freedom of the colloids are constrained to be along the local tangent vector of the filament and are hence not true degrees-of-freedom of the system [23]. The filament boundary conditions are clamped at the base and free at the tip, such that the equilibrium shape is a straight filament (Fig. 1C).

Of the family of models possible within the above active filament framework, we considered the simplest one where activity is confined to only the distal colloid and is due to a potential dipole of strength *d*_0_: the simplest hydrodynamic representation of a finite-sized, force-free swimmer. This “swimmer on a string” is a much simplified model for the ciliary forcing in organisms like *L. olor*, but more generally can be considered as the simplest model for morphodynamics where there is an interplay between active stresses exerted on an ambient fluid and material response of the cell or soft-robot body. Importantly, the orientation of this potential dipole activity is constrained to be along the local tangent at the filament tip. This so-called tip “follower-force” is inspired by ciliates like *L. olor* wherein the cilia are anchored to the cell membrane and underlying cytoskeleton and thus their beat-plane, and hence thrust direction is constrained by the local orientation of the organism. While the angle between the active stresses and the filament tangent can have non-zero values and is indeed dynamic in *L. olor*, we consider the simplest case of where the stresses are tangential at all times and only switch direction to model ciliary reversals.

To model the dynamics of the active stresses of the tip colloid models we considered a deterministic activity profile such that the sign of the potential dipole changes sign periodically at a fixed time scale *τ*_*a*_, switching between phases of compression and extension of the filament (Fig. 1C). This leads to a square-wave profile of the activity strength of the tip colloid given by:

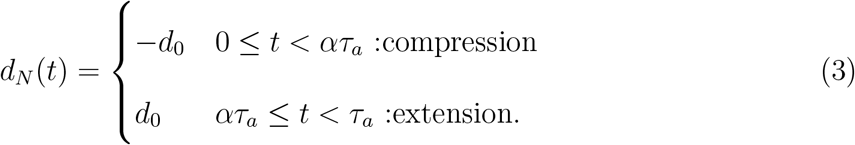

Here 0 ≤ *α* ≤ 1 is the duty-cycle corresponding to the duration of the compressive phase relative to the total activity time scale. We chose *α* = 0.5 for all our simulations leading to equal duration of the compressive and extensional phases. The square-wave activity profile chosen has a biophysical basis in the brief transitory phase between the much longer forward and reversed beat phases of the cilia [7].

Our active filament model thus consists of *N* colloids, with the distal colloid being active while the remaining *N* − 1 are passive, giving a filament of equilibrium length *L* = (*N* − 1)*b*_0_. The system can be made dimensionless using the colloid radius *a* as the length scale, the stretch relaxation time scale *µL/k*, velocity scale *a*^2^*d*_0_ and the force and moment scales of *µa*^3^*d*_0_ and *µa*^3^*d*_0_*L*, respectively. We can also define a dimensionless activity strength which is the ratio of active and elastic restoring moments given by: 𝒜 = *µa*^3^*d*_0_*L*^2^*/κ* [17]. All quantities in the rest of this work are presented in dimensionless terms unless otherwise indicated.

We validated our model using two case studies: one static and one dynamic (Supplementary Information §1.2). In the first we compared the steady state shape of a filament under a fixed transverse force to theoretical predictions for a Euler-Bernoulli beam and find good agreement (Supplementary Information §1.2). In the second we simulated the filament dynamics under a constant compressive tip follower-force and compared the resulting frequency with the scaling law predicted by [15], and again find good agreement with the predicted −4*/*3 power-law (Supplementary Information §1.2). This “flapping” time scale can be regarded as a natural active time scale of the system for a given set of filament parameters, and for our system can be written as:

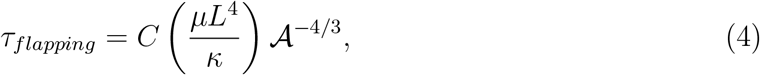

where *µL*^4^*/κ* is the equilibrium time scale for the relaxation of bending perturbations of the filament, and *C* is an 𝒪(1) constant.

### 2.2 Periodically driven active filaments display complex behaviors

To explore the space of filament behaviors, we evolve the dynamics of filaments starting from various straight initial shapes, over hundreds of activity cycles, with a deterministic activity profile (given by Eq. (3)), and for a range of activity strengths *A* and times *τ*_*a*_. For weak activity strengths, or short activity times, there is no buckling during the compressive phase, and the filament remains straight and the only dynamics is a small change in filament length in phase with the driving signal. For higher activity strengths and durations, the filament dynamics during an activity cycle follows a typical pattern wherein the compressive phase causes the filament to buckle in-plane and the tip of the filament to decorrelate in orientation and also rapidly change position (Fig. 2A). Note that this buckling is not the classical Euler buckling of an elastic beam under constant load, but is due to the so-called tip “follower-force” since the direction of compressive force is determined by the tip orientation itself (Fig. 1C, Fig. 2B, C) [16, 17]. This is made apparent by visualizing the flow around the filament during the compressive and extensional phases, with a strong coupling observed between the filament geometry and the active flow and hence, stress generated by the tip (Fig. 2B, C; Movie S1). The subsequent extensional phase largely preserves the tip orientation, but the filament as a whole now points in a new direction (Fig. 2A). As the filament extends, the “swimmer” reaches the end of its tether leading to lower tip speeds and filament displacements during the extensional phase, but faster flows around the filament (Fig. 2A, C). Each cycle of compression and extension thus leads to a change in filament shape and orientation. With regards to the filament tip, which one can think of as the main driver of the filament dynamics, the filament behavior for high enough activity strength resembles a rapid orientation decorrelation, followed by correlated movement of the tip analogous to the tumbles and runs in free-swimming cell motility [28](Fig. S3).

**Fig. 2:**
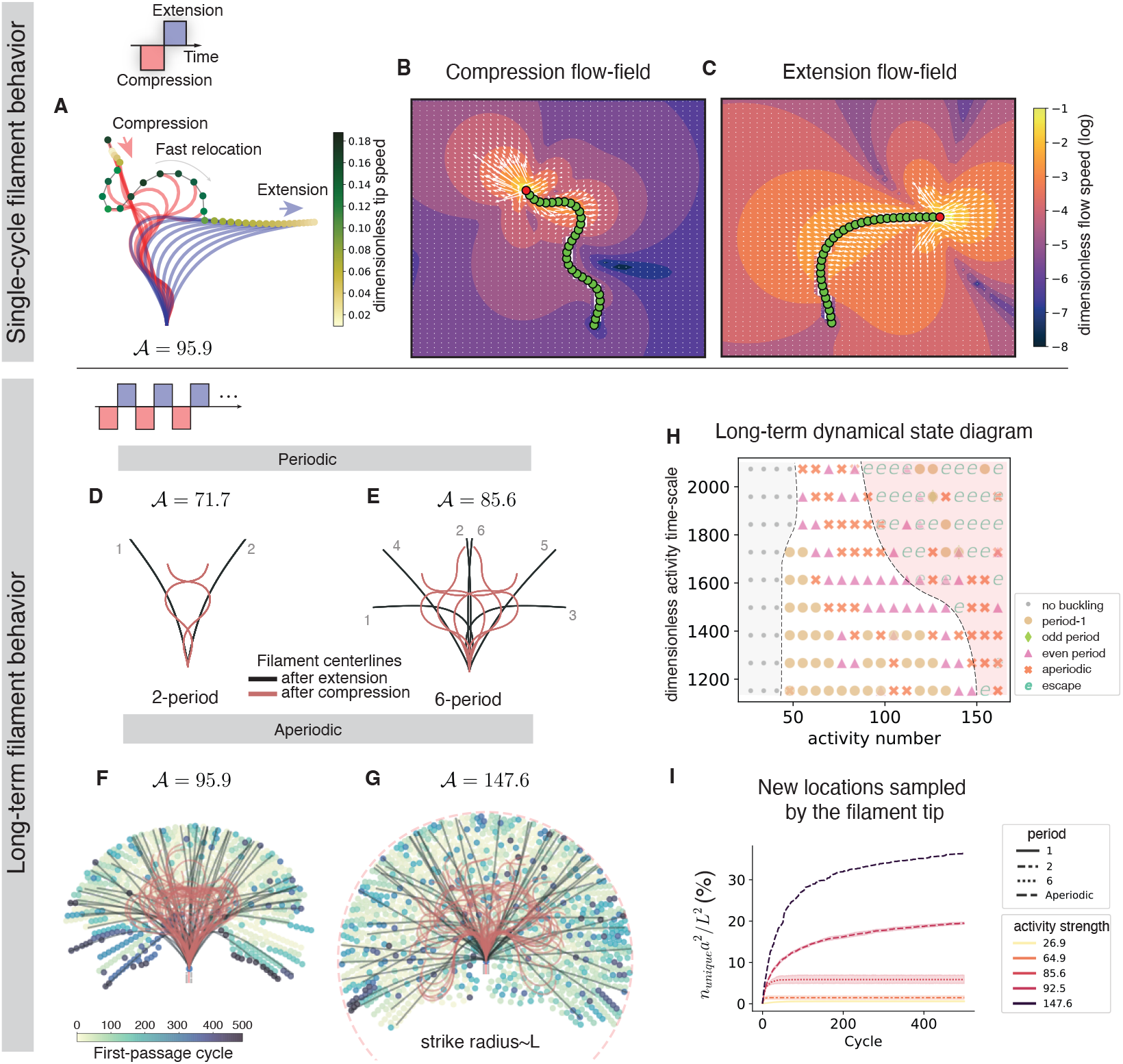
Emergent behavior in active filaments under dynamic follower-forces. **(A)** Filament center-line shape dynamics during a single cycle of compression and extension. The filament center-lines are colored by activity phase (red: compression, blue:extension). Filament tips are colored by the dimensionless tip speed. **(B)** and **(C)** The instantaneous flow around the filament during compression and extension phases, respectively, with color denoting the dimensionless flow speed (log-scale). **(D)-(G)** Long-time filament dynamics (over 500 activity cycles) for varying activity strengths (𝒜) and *τ*_*a*_ = 1744, showing both periodic (D, E) and aperiodic behavior (F, G). Filament center-lines shown at the end of compression (red) and extension (black) phases for the last 100 activity cycles. In (F, G) the unique locations sampled by the filament tip are shown colored by the first-passage cycle (points are not to scale). The activity strength is given by 𝒜 = *µa*^3^*L*^2^*d*_0_*/κ*, where *µ* is the fluid viscosity, *a* the filament/colloid radius, *L* the filament rest-length, *d*_0_ the potential-dipole strength of the tip colloid and *κ* the bending rigidity. **(H)** Long-term filament behavioral state diagram for varying activity strength and time scale over 500 activity cycles. The dashed lines and shaded regions are a guide to the eye. Each symbol represents simulations from one initial condition (3 initial conditions per parameter value pair: a total of 540 simulations run over 500 cycles). **(I)** Number of new locations visited by the filament tip (*n*_*unique*_) as a function of activity cycles, reported as percent of the maximum number of accessible locations to within a filament diameter, which scales as *L*^2^*/a*^2^, where *a* is the filament radius and *L* is the filament length. The lines represent the mean and the shaded regions the 95% confidence-interval over 7 initial conditions.

Over many iterative activity cycles, we find a rich space of filament behaviors (Fig. 2D-H). We observe period-doubling dynamics where the filament shapes repeat every 2, 4, 6 … cycles, as well as period-1 cycles where the filament repeatedly samples a particular angular location away from the symmetry axis (Fig. 2D, E, H; Movie S2). Intriguingly, above a critical activity strength we observe aperiodic dynamics where the filament shapes never repeat even for long times (Fig. 2F, G, H; Movie S2). The sequence of filament shapes appears unpredictable and random for the aperiodic cases: for instance, even the half-plane that the filament tip occupies after each cycle (left vs right) appears to be a random sequence (Movie S3). At higher activity strengths we also observe that filaments can “escape” from the half-space, which we define as when the angle between the first two colloids at the base and the equilibrium angle becomes greater than *π/*2 (Fig. 2H).

The aperiodic filament dynamics also has an important consequence for how the tip of the filament samples space, a function of interest for biological systems [29, 30, 31, 32]. In striking contrast to periodic dynamics where the filament tip samples only a small number of locations repeatedly, aperiodic dynamics leads to the tip sampling up to ≈ 35% of the maximum available unique locations near the filament over 500 cycles (Fig. 2I). Further, the tip exploration of space is not systematic in nature, but appears “stochastic” with the tip darting between points on either side of the symmetry axis. This is further observed in the well-mixed nature of the spatial distribution of first-passage times for each unique location visited by the tip (defined as locations separated by more than twice the colloid radius) (Fig. 2F, G). Thus even periodic deterministic activity patterns generate aperiodic, “search-like” behaviors in this simple configuration of active filaments with tip follower-forces.

Overall even when driven by deterministic activity patterns, the combination of compressive and extensional follower-forces leads to a very rich behavioral space (Fig. 2H). Next we characterize these behaviors further using tools from non-linear dynamics to explain their emergence.

### 2.3 A transition to chaos underlies aperiodic filament dynamics and dense spatial sampling

Are the observed aperiodic behaviors a signature of chaotic dynamics? To answer this question we created a reduced-order representation of the filament dynamics using Principal-Component-Analysis (PCA). We computed the eigenvectors corresponding to the covariance matrix of the tangent angle representation of the filament center-lines *θ*_*t*_(*s, t*), where *θ*_*t*_ is the local tangent angle at the arc length location *s* and at time *t*, drawn from simulations over 500 activity cycles. By projecting the filament shapes onto these eigenmodes, we found the time-varying amplitudes that describe filament dynamics. For low and moderate activity strengths (30 ≤𝒜 ≤ 100, and *τ*_*a*_= 1744), we find that the first 3 eigenmodes are sufficient to explain 95% of the filament shape variance (Fig. 3F). Using this representation, the filament dynamics for low and moderate activity becomes a trajectory in an ℝ^3^ “shape-space”, with the origin corresponding to the equilibrium, straight-filament shape (Fig. 3C, D).

**Fig. 3:**
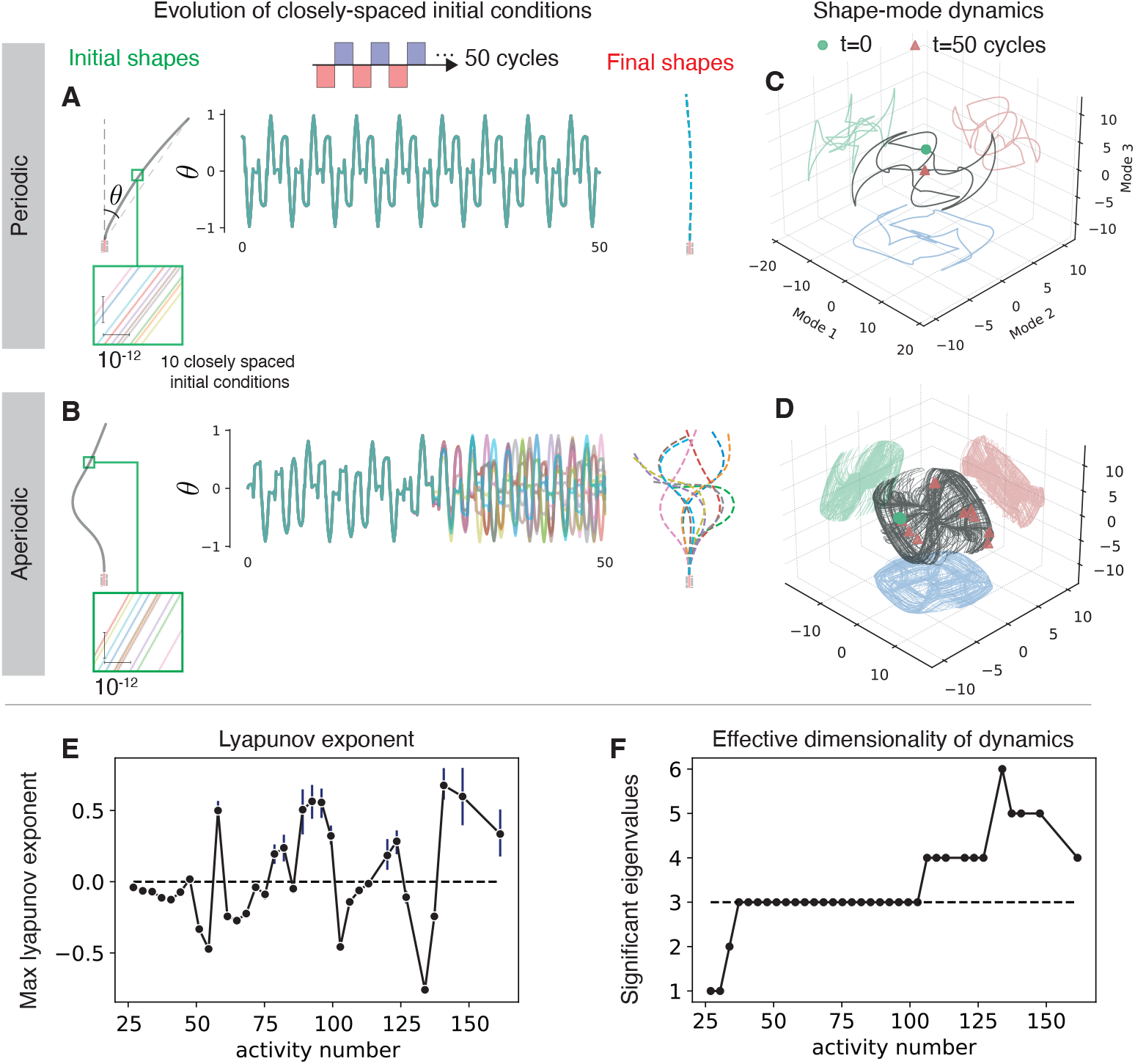
A transition to chaos underlies aperiodic filament behavior. **(A)** and **(B)** Evolution of 10 closely spaced initial filament shapes (pairwise euclidean distance *<*= 10^*−*12^), driven by deterministic activity profiles. Initial and final shapes are plotted, along with time-evolution of the filament base-tip angle. For the periodic case (A), the final shapes all lie on top of each other and are indistinguishable at the scale of the filament radius. In striking contrast, for the aperiodic case (B), closely spaced filament shapes evolve into distinct final shapes over just 50 cycles. **(C), (D)** Visualization of filament shape dynamics using a lower-dimensional representation by projecting onto an orthogonal set of eigenvectors (shape modes). For both periodic and aperiodic cases the filament dynamics settles onto an attractor of finite size (black trajectory). The projections of this trajectory along the coordinate axes are shown as red, green and blue curves. (C) Periodic dynamics results in closed-loops in shape space, while (D) aperiodic dynamics leads to trajectories that densely sample space but remain bounded. (C),(D) Evolution of closely spaced initial conditions (green dots) into final shapes (red triangles) for the periodic and aperiodic cases are overlaid. **(E)** Maximum Lyapunov exponents as a function of activity strength calculated from the growth/decay rate of small perturbations. Positive values are a feature of chaotic dynamics, while negative values correspond to periodic dynamics. Exponents are calculated over 100 unique initial conditions chosen on the attractor, for each of which we evolved 10 closely-spaced initial shapes. **(F)** Number of eigenvalues needed to explain 95% of the shape variance as a function of the activity strength. The number of active shape modes is a measure of the dimensionality of the dynamics.

We find that periodic dynamics leads to a limit cycle in shape-space, while aperiodic dynamics leads to flow on a much more complex, but finite-sized attractor (Fig. 3C, D). To test if the dynamics was chaotic, we chose 100 random points along this attractor and seeded 10 nearby initial conditions by adding small transverse perturbations of amplitude *δ*_0_ = 10^*−*12^ to the filament shape at that point. We then computed the pairwise Euclidean distance between filament shapes evolved from these initial conditions. The pairwise Euclidean dis-tance between filament shape *m* and *n* is defined as 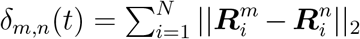 where 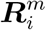 is the position of the *i*^*th*^ colloid on filament *m*. We find that the pairwise distances grow exponentially for aperiodic dynamics, leading to the nearby trajectories ending up spread across the attractor and leading to distinct filament shapes even over 50 cycles (Fig. 3B, D; Movie S5). In contrast periodic dynamics does not lead to exponential growth, and nearby trajectories remain close (to within 1% of the filament radius) even for long times (Fig. 3A, C; Movie S4).

We further estimated the growth or decay rate of these perturbations or the maximum Lyapunov exponent by fitting a function of the form 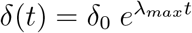 to the pairwise distance trajectories starting at 100 initial locations spread across the attractor. Calculated for varying activity strengths, we find that at low enough activity strengths (𝒜 < 50) the dynamics is always periodic corresponding to *λ*_*max*_ *<* 0 (Fig. 3E). We also find that beyond a critical activity we can have *λ*_*max*_ *>* 0, corresponding to aperiodicity (Fig. 3E). Interestingly, we also find patches of negative growth rates (*λ*_*max*_ *<* 0) for higher activities corresponding to windows of periodic dynamics (for instance see the window around 𝒜 ≈ 100 in Fig. 3E). Based on the typical values of computed positive growth rates, we find that practically indistinguishable initial filament shapes (to within 10^*−*12^ in pairwise distance) become completely distinct over *O*(50) activity cycles (Fig. 3B, D, E; Movie S5).

Overall, we find that the filament dynamics above a critical activity strength is aperiodic for long times, deterministic, and highly sensitive to initial conditions (positive Lyapunov exponent), thus satisfying all the conditions for chaos [33]. This chaotic dynamics underlies the unpredictable sampling behavior observed for the filament (Fig. 2D, E), and also leads the filament to sample ever newer shapes and tip locations, thus minimizing spatial resampling. The above discovery of chaotic dynamics places the behavior of *L. olor* in a new light, since exploiting these dynamics offers unique advantages in rapidly sampling the extracellular environment and explains the highly dynamic shape changes of the neck. At the same time chaotic dynamics will place strong constraints on the control of these behaviors in a biological context where noise is an important factor. To explore control of filament dynamics using dynamic activity and possible design strategies for programming filament behaviors, we now explore the fundamental non-linearities inherent in the follower-force compression and extension operation that underlie periodic and chaotic long-term dynamics.

### 2.4 A 1D map of filament shape to predict and design long-term filament behavior

How do periodic and chaotic behaviors emerge from the underlying active filament system and dynamic follower-force activity profiles? And how do the emergent long-term behaviors depend on the underlying properties of the filament and activity dynamics? We next present an analysis that seeks to explain the observed behaviors by exploring how non-linearities arise in this simple system and how they are connected to underlying system parameters.

Overall our strategy is to postulate that the filament dynamics can be approximated by a simple map acting iteratively on the filament orientation angle, and to prove this postulate *a posteriori*. We then analyze the fixed points of this map (which correspond to periodic, limit cycle filament dynamics) and their linear stability to predict filament behaviors from the shape of the map. We now describe our analysis in further detail.

We interpreted the dynamics as a discrete iterated map operating on the filament shape (Fig. 4 A). We considered the simplest version of this map that involves only the filament orientation angle *θ*_*n*_, defined as the angle between a ray connecting the base of the filament to its tip and the filament’s equilibrium orientation, after *n* activity cycles (Fig. 4A). We plotted this map (i.e. *θ*_*n*+1_ vs *θ*_*n*_) using data from full numerical simulations and observed that it showed a lot of structure for the case of aperiodic filament dynamics, underscoring the the fact that these dynamics are chaotic but not stochastic (Fig. 4B, C) [34]. Next we computed the qualitative shape and features of this map, approximated as a function, *f* (*θ*), where *θ*_1_ = *f* (*θ*_0_), by simulating a single cycle of compression and extension and computing the final angles *θ*_1_ given a range of initial angles *θ*_0_ ∈ [0, *π/*2]. Note that this does not approximate the underlying dynamics, but assumes the filament is straight, and pre-stretched (corresponding to an extensional cycle at the appropriate activity strength and time scale) at the beginning of the simulated activity cycle. We found that, for both varying activity strength and time scale, the predicted *f*(*θ*) closely matches the overall shape and qualitative features of the map from full simulations over many hundreds of cycles (Fig. 4B, C).

**Fig. 4:**
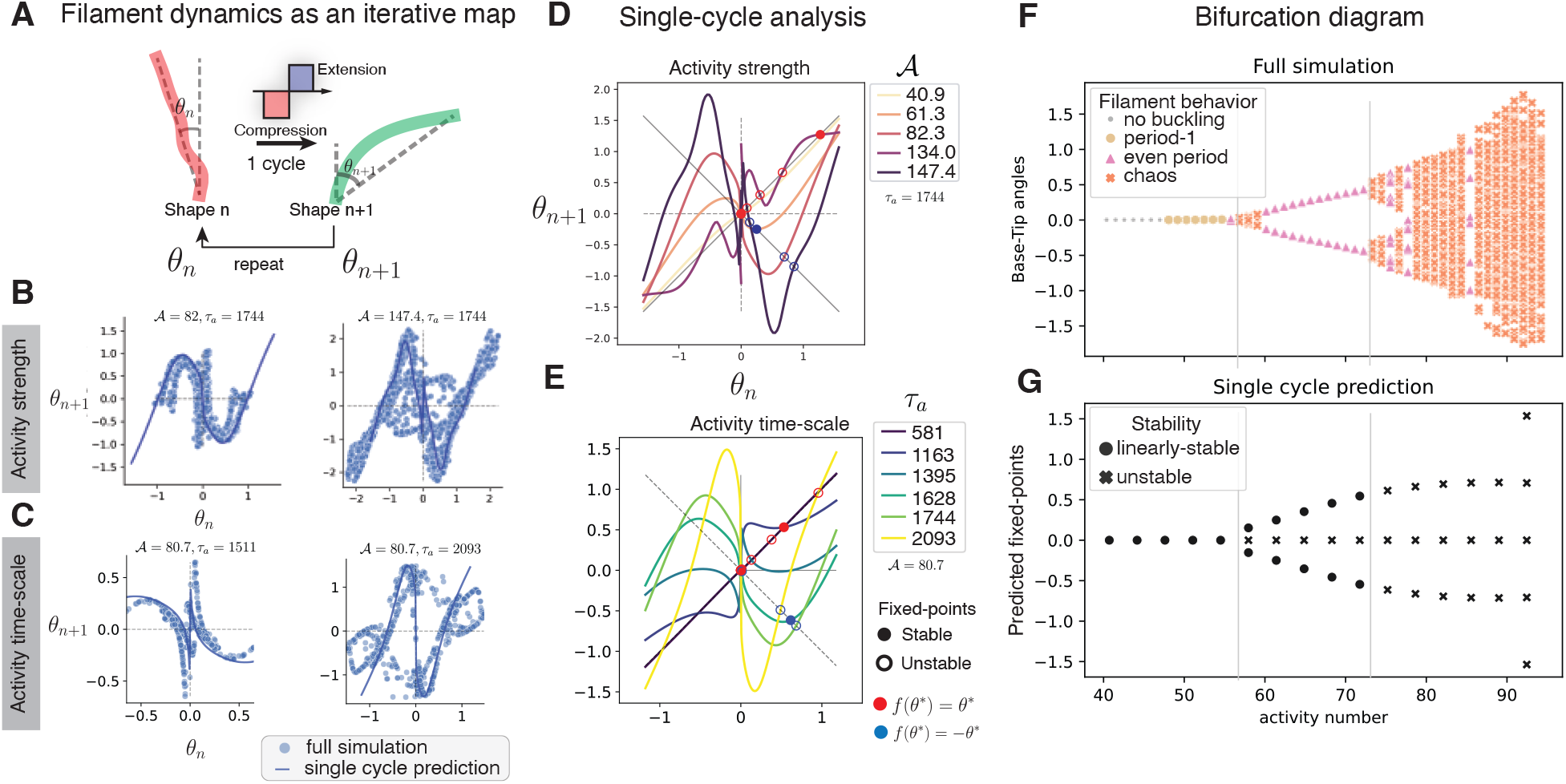
A non-linear 1D map predicts long-term filament behavior. **(A)** Filament dynamics during an activity cycle (compression+extension) as a 1D iterated map of the filament base-tip angle *θ*. Long-term filament dynamics can then be regarded as iterations of this map, i.e. *θ*_*n*+1_ = *f* (*θ*_*n*_), where *n* is the activity cycle number. **(B), (C)** Iterated maps of filament angle for different activity strengths (B) and time scales (C). The dots correspond to full numerical simulations, and the solid lines to the predicted map over a single-cycle. **(D), (E)** The shape of the map *f* (*θ*) from the single-cycle prediction plotted for varying activity strength (D) and time scale (E). The fixed points of *f* (*θ*) = *θ* and *f* (*θ*) = −*θ*, and their stability, correspond to the existence and the stability of period-1 and even period limit cycles, respectively, and hence predict the long-term dynamics of the filament. Filled and empty circles correspond to stable and unstable fixed points, respectively. The red and blue circles correspond to fixed points of *f* (*θ*) = *θ* and *f* (*θ*) = −*θ*, respectively. **(F)** Bifurcation diagram based on the filament base-tip angles sampled at the end of the extension phase, for full numerical simulations. With increasing activity strength successive period-doubling bifurcations eventually lead to chaos. Symbols correspond to the nature of the dynamics. **(G)** Predicted bifurcation diagram from the single-cycle analysis. Filled circles: stable fixed points, crosses: unstable fixed points. Note that the single cycle analysis predicts the onset of the first period-doubling behavior as well as the onset of chaos (vertical gray lines).

Next we analyzed two types of fixed points of the map *f*(*θ*) as well as their stability to predict long-term filament behavior. Firstly, we find that due to symmetry of the system about *θ* = 0, *f* (*θ*) must be odd, i.e. *f*(−*θ*) = −*f*(*θ*). This implies that the fixed points *θ*^∗^ of *f* (*θ*) = −*θ* are also of interest since we can show that they are also fixed points of any even-iterate map *f*^2*m*^(*θ*), where *n* ∈ ℕ (Supplementary Information §1.3), which corresponds to 2*n*-period limit cycles of the filament dynamics. The stability of both types of fixed points can be evaluated by the eigenvalue that governs the growth rate of perturbations about that fixed point/ limit cycle, given by *λ* = |*f*′(*θ*^*∗*^)|, where the ′ signifies a derivative. The fixed point is linearly stable/unstable for *λ <* 1 and *>* 1, respectively (Supplementary Information §1.3). The fixed points and their stability for the single-cycle iterated maps computed for varying activity strength and time scale are shown in Fig. 4D and E, respectively.

Using this stability analysis we predicted the long-term filament dynamics. We visualized the dynamics as a bifurcation diagram in the observed filament orientation angles *θ*_*n*_ at the end of each extension phase as a function of activity strength and found good agreement with the fixed-points and their stability predicted using the single-cycle analysis above(Fig. 4F, G). As seen in Fig. 4G, at low activity, we predict no buckling, followed by dynamic-equilibrium in the form of period-1 dynamics where the origin (straight filament) remains a fixed point. At higher activity strengths, we see the first period-doubling bifurcation and eventually the onset of chaos, which is due to a cascade of period-doubling bifurcations (Fig. 4F). The single-cycle prediction closely matches the observed onset of the first period-doubling bifurcation, as well as the onset of chaotic dynamics (Fig. 4F, G).

The shape and fixed points of the single-cycle iterated map allows us to infer, at a glance, how long-term filament dynamics emerges due to periodic follower-force actuation, making these curves an important design tool. At low activity strengths and/or activity time scales, the map is almost diagonal, since the activity is too weak or short-lived to effect a large change in the filament orientation (Fig. 4D, E). With increasing activity strength/time scale, the map deviates from the *f*(*θ*) = *θ* diagonal, but is still nearly linear, with the origin as the only, stable, fixed point, which corresponds to a period-1 cycle (Fig. 4F, period-1 cycles). With further increase in activity strength or time scale, we see the first bifurcation to a period-2 dynamics happen as the map develops a valley and intersects the negative diagonal at a stable fixed point, since *f*′(*θ*^*∗*^) *<* 1 (Fig. 4D, E). The location of this fixed point also gives us the angular location that the filament tip repeatedly returns to (Fig. 4F, G, period-2m cycles). With further increase in activity strength or time scale, the valley in the map becomes steeper such that the fixed point of the even iterate map eventually loses stability when |*f*′(*θ*^*∗*^)| *>* 1 via a flip-bifurcation (Fig. 4D, E) [33]. Interestingly, as shown above, this implies that all even iterate maps *f*^2*m*^(*θ*), and hence, all period-2m cycles, are now unstable. This then is the transition point to chaotic dynamics since all even periodic cycles are now unstable (Fig. 4F). The analysis above allows us a ready means, by analyzing the two major classes of fixed points, to infer long-term dynamics from the shape of *f*(*θ*). A more detailed analysis will be necessary to predict the existence of other interesting features observed in our simulations such as periodic windows within the chaotic regions of parameter space (Fig. 4F), transient chaos, as well as exotic odd-periodic cycles (Fig. 2H).

We now use insights from these predicted non-linear maps to explore how filament behaviors may be programmed by modulated activity signals.

### 2.5 Programming filament behavior using modulated follower-force activity

Here we briefly demonstrate how functional filament behaviors can be programmed through modulated activity signals by combining results from the non-linear dynamics of follower force compression and extension (Fig. 4). These examples serve as a first step in understanding the space of possible behaviors in a biologically accessible parameter space including amplitude and frequency modulation of the activity signals as well as the effects of noise in the signals.

A useful control capability for active filaments is the ability to quickly return the filament to its equilibrium orientation. We show that such a shape “reset” can be performed using activity by modulating the activity pulses that drive the filament. We use the return-map profiles in Fig. 4D, E as a design tool, and leverage the existence of a “dynamic equilibrium” of the straight filament state as a fixed point of the dynamics at low activity strengths or time scales (Fig. 4D, E). Thus by driving the filament at a low activity strength level or fast time scale we expect to drive the filament to a straight shape, at a much faster rate than due to equilibrium bending forces (Fig. 5A). We confirmed this using simulations starting from 100 randomly chosen filament shapes and showed that these could be quickly driven close to the straight filament shape in ≈ 50 cycles for frequency modulation and even faster in ≈ 10 cycles for amplitude modulation (Fig. 5A, Movie S6). In comparison the equilibrium return to the straight filament would take a time *t* ∼ *µL*^4^*/κ*, equivalent to ≈ 300 cycles at the activity time scale considered.

**Fig. 5:**
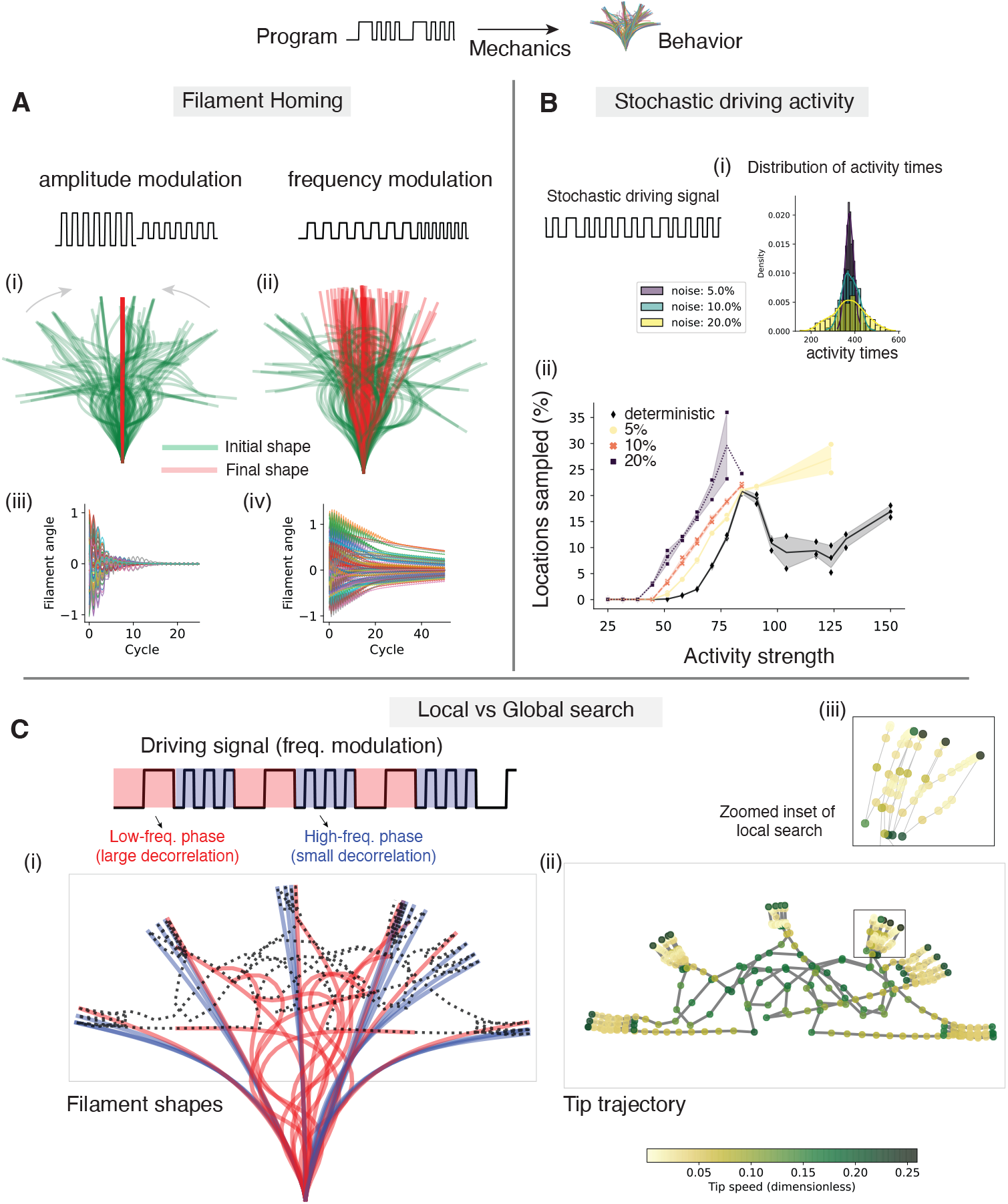
Programming filament behavior using modulated activity signals and exploiting non-linear follower-force dynamics. **(A)** Demonstration of filament homing using (i) amplitude or (ii) frequency modulated activity signals. Overlay of filament center-lines at the start (green) and end (red) of 50 cycles of activity starting from 100 random filament shapes. Time evolution of the filament base-tip angle for (iii) amplitude and (iv) frequency modulation cases. **(B)** Effects of stochasticity/noise in the durations of the compressive and extensile phases of the activity signal. (i) The distribution are plotted for increasing values of noise-strength (variance). (ii) Unique locations sampled by filament tip, as a percent of maximum, for the deterministic and stochastic profiles with increasing noise strength. Individual data points for 3 initial conditions are shown and the shaded area represents a 95% confidence interval. **(C)** Programming local and global sampling of the filament tip using alternating periods of low and high frequency activity. (i) Filament shapes during low and high frequency states shown in red and blue, respectively, with the tip trajectory (dashed-line). (ii) Plotting the tip speed reveals that the filament dwells in a local region during high-frequency states and rapidly moves across space during the low-frequency state. (iii) Zoomed in view of a local sampling trajectory.

While so far we have considered the effects of a periodic activity profile to elucidate the underlying dynamics of the follower-force filament system, we next explored how noise or stochasticity in the durations of the driving signal affects the emergent behaviors (Fig. 5B). We find that Gaussian noise, in general, increases the sampling efficiency of the filament (Fig. 5B). For example, with strong noise (*σ*_*a*_*/*⟨*τ*_*a*_⟩ = 20%) the unique locations sampled is ≈ 35% of the maximum compared to 25% for a purely deterministic driving signal (Fig. 5B). Overall, these results show that stochasticity in the driving signals alone cannot lead to effective sampling of filament shapes at these non-Brownian length scales, and that effective tip exploration of space relies on the underlying non-linear and chaotic filament dynamics.

Finally, we show that more complex filament behaviors can be programmed by combining sequences of activity pulses. Specifically we show how one can use alternating periods of low and high-frequency activity pulses to perform global and local sampling of space by the filament tip (Fig. 5C, Movie S6). Low-frequency pulses drive follower-force bending in the non-linear regime wherein there is a large change in the filament orientation thus transporting the tip to a new location in space. On the other hand, high frequency pulses drive the filament dynamics in the linear regime, causing the filament orientation to be mostly correlated and the tip to explore a small cone of space near the current filament orientation (Fig. 5C, Movie S6). We also find that the tip speeds are lower during the high-frequency phase and higher during the low-frequency phase (Fig. 5C). Overall such activity profiles allow us to implement the equivalent of an “intermittent search” strategy wherein the filament tip has a higher chance of locating a target when searching locally, due to the slower tip speeds, but can then quickly relocate to a new region of space to minimize resampling.

## 3 Discussion and Outlook

In this work, we have described an active filament model driven by periodic phases of extensional and compressive tip follower forces and characterized the rich space of behaviors including the discovery of chaotic dynamics. This dynamical behavior is novel and is in striking contrast to other active elastohydrodynamic systems such as cilia which are typically characterized by periodic limit cycle dynamics [35]. Unlike earlier active filament models [15] that need to appeal to noise to achieve aperiodicity and mixing, our work points to a simple activity motif that results in chaos even at non-Brownian length scales, thus significantly expanding the functional space of active filaments - from molecules to cellular protrusions to microscale robotics. By generalizing the driving activity to have time-varying phases of extension and compression, we also introduced a fundamental notion of programmability, wherein a “program” (time sequence of activity) is directly converted to “behavior” (dynamical states) through a non-linear mapping function which can be derived from first-principles of non-variational mechanics.

Our work also indicates that the non-linear, chaotic dynamics of a filament under a time-varying follower-force is likely an important underlying mechanism through which *L. olor* achieves its remarkable behaviors that include dense and rapid sampling of space around the cell using its active slender protrusion. To the best of our knowledge, our work constitutes the first example of the utility of chaos in the behavior of a single-celled organism. In biological implementations of this framework such as in *L. olor*, this intrinsic mechanical non-linearity is likely amplified further by other effects such as hysteric variations in the material property of the neck, twist-buckling due to torsional activity, and noise in the periods of the forward and reversed ciliary beat phases. Active filament models with dynamic activity could also serve as general frameworks for understanding behavior in other ciliates where force generators (cilia) are anchored directly to cell membranes and the cytoskeleton, whose geometry is in turn a function of active stresses. These cilia-covered cell surfaces [13] are a hallmark of one of the largest group of protists, and our work is a first step towards a general biophysical framework for the rich class of behaviors observed in these organisms.

More generally, our work indicates that periods of compression and extensional activity are a novel solution to the problem of how a microscale tethered object can achieve orientation decorrelation and search its surrounding space - a classical problem in mathematics [30, 36]. This solution allows an organism like *L. olor* or a microscale-robot to search the space around itself. In this way, active filaments with time-varying follower-force activity are a simple, mechanical embodiment of “intermittent search behavior” in unicellular organisms like *L. olor*, a behavior that is typically observed in higher animals possessing nervous systems [31, 32, 29, 37]. Further, while we calculated search metrics using the filament radius as the size scale, the hydrodynamics leading to a fluid flux towards the filament tip during the slow extensional phase allows it to “scan” an effective volume of fluid which is much larger than that due to the filament radius, hence significantly expanding the search space (see Fig. 2C). Lastly, while we have looked at search behavior in an unbounded fluid, it is likely that this type of emergent spatial sampling which exploits non-linearity and chaos has significant advantages in more complex, crowded environments such as porous media, where deterministic strategies might fail due to the presence of obstacles [38].

Our work is a first step towards understanding the programmability of the state space of active filament behaviors under dynamic follower-forces and the attendant consequences for both engineering and biological applications. The simplicity of the proposed active filament system and periodic driving signals suggests the possibility of experimental realizations (at both macro and micro-scales) and theory-experiment dialogue to fully map out the space of filament behaviors. From an engineering context, our work suggests a new class of self-driven, soft, microscale robotic arms that can leverage the non-linear dynamics and chaotic motions discovered here to achieve novel functional behaviors. Lastly, from a biological perspective, it will be interesting to explore how sensing and control strategies have evolved to shape behavior under the constraints of self-driven active filament dynamics discovered here.

## Authors’ Contributions

D.K., M.P designed the research; D.K developed the *in silico* model, wrote the code and carried out simulations. D.K. carried out the data analysis. D.K. and M.P. wrote the paper.

## Data Availability

Simulation data will be made available in the Dryad data repository (https://doi.org/10.6078/D12T52). The code used for simulations, analysis and figure generation can be accessed at the following GitHub repository: (https://github.com/deepakkrishnamurthy/PyFilaments).

## Acknowledgements

We thank Scott Coyle and Christina Hueschen for discussions and comments. We thank Ellie Flaum, Ray Chang, Samhita Banvar and all members of the Prakash lab for discussions. D.K acknowledges support from Bio-X fellowship and the Schmidt Science Fellowship. M.P. acknowledges financial support from NSF Career Award, Moore Foundation, HHMI Faculty Fellows program, NSF CCC (DBI-1548297) program, NSF Convergence Award (OCE-2049386), Schmidt Foundation and CZ BioHub Investigators program.

## Notes

https://doi.org/10.6078/D12T52

### Competing Interest Statement

The authors have declared no competing interest.

## References

[1] Brown, A. E. X. & de Bivort, B. Ethology as a physical science. Nat. Phys. 14, 653–657 (2018).

[2] Sponberg, S. The emergent physics of animal locomotion. Phys. Today 70, 34–40 (2017).

[3] Stephens, G. J., Johnson-Kerner, B., Bialek, W. & Ryu, W. S. Dimensionality and dynamics in the behavior of c. elegans. PLoS Comput. Biol. 4 (2008).

[4] Jordan, D., Kuehn, S., Katifori, E. & Leibler, S. Behavioral diversity in microbes and low-dimensional phenotypic spaces. Proc. Natl. Acad. Sci. U. S. A. 110, 14018–14023 (2013).

[5] Berman, G. J., Bialek, W. & Shaevitz, J. W. Predictability and hierarchy in drosophila behavior. Proc. Natl. Acad. Sci. U. S. A. 113, 11943–11948 (2016).

[6] Jennings, H. S. Behavior of the Lower Organisms (Columbia University Press, 1906).

[7] Coyle, S. M., Flaum, E. M., Li, H., Krishnamurthy, D. & Prakash, M. Coupled active systems encode an emergent hunting behavior in the unicellular predator lacrymaria olor. Curr. Biol. 29, 3838–3850.e3 (2019).

[8] Coyle, S. M. Ciliate behavior: blueprints for dynamic cell biology and microscale robotics. Mol. Biol. Cell 31, 2415–2420 (2020).

[9] Wan, K. Y. & Jékely, G. Origins of eukaryotic excitability. Philos. Trans. R. Soc. Lond. B Biol. Sci. 376, 20190758 (2021).

[10] Bull, M. S. & Prakash, M. Mobile defects born from an energy cascade shape the locomotive behavior of a headless animal (2021). 2107.02940.

[11] Yanase, R., Nishigami, Y., Ichikawa, M., Yoshihisa, T. & Sonobe, S. The neck deformation of lacrymaria olor depending upon cell states. Journal of Protistology 51, 1–6 (2018).

[12] Miller, S. The predatory behavior ofdileptus anser. J. Protozool. 15, 313–319 (1968).

[13] Lynn, D. The Ciliated Protozoa: Characterization, Classification, and Guide to the Literature (Springer, 2008).

[14] Wood, W. & Martin, P. Structures in focus–filopodia. Int. J. Biochem. Cell Biol. 34, 726–730 (2002).

[15] Chelakkot, R., Gopinath, A., Mahadevan, L. & Hagan, M. F. Flagellar dynamics of a connected chain of active, polar, brownian particles. J. R. Soc. Interface 11 (2014).

[16] Laskar, A. & Adhikari, R. Filament actuation by an active colloid at low reynolds number. New J. Phys. 19 (2017).

[17] De Canio, G., Lauga, E. & Goldstein, R. E. Spontaneous oscillations of elastic filaments induced by molecular motors. J. R. Soc. Interface 14 (2017).

[18] Bolotin, V. V. Nonconservative problems of the theory of elastic stability (Macmillan, 1963).

[19] Ling, F., Guo, H. & Kanso, E. Instability-driven oscillations of elastic microfilaments. J. R. Soc. Interface 15, 20180594 (2018).

[20] Banavar, S., Flaum, E. & Prakash, M. Hydraulics of cellular extension and contraction in lacrymaria olor. In APS March Meeting 2022 (American Physical Society, 2022).

[21] Brette, R. Integrative neuroscience of paramecium, a “swimming neuron”. eNeuro (2021).

[22] Marr, D. Vision: A computational investigation into the human representation and processing of visual information. The MIT Press (MIT Press, London, England, 2010).

[23] Laskar, A. & Adhikari, R. Brownian microhydrodynamics of active filaments. Soft Matter 11, 9073–9085 (2015).

[24] Singh, R. & Adhikari, R. Generalized stokes laws for active colloids and their applications (2016).

[25] Singh, R. & Adhikari, R. PyStokes: phoresis and stokesian hydrodynamics in python. Journal of Open Source Software 5, 2318 (2020).

[26] Guazzelli, É. & Morris, J. F. A Physical Introduction to Suspension Dynamics (Cambridge University Press, 2011).

[27] Singh, R., Ghose, S. & Adhikari, R. Many-body microhydrodynamics of colloidal particles with active boundary layers. J. Stat. Mech: Theory Exp. 2015 (2015).

[28] Berg, H. Random Walks in Biology (New and Expanded Edition).

[29] Bénichou, O., Loverdo, C., Moreau, M. & Voituriez, R. Intermittent search strategies. Rev. Mod. Phys. 83, 81–129 (2011).

[30] Holy, T. E. & Leibler, S. Dynamic instability of microtubules as an efficient way to search in space. Proc. Natl. Acad. Sci. U. S. A. 91, 5682–5685 (1994).

[31] Viswanathan, G. M. et al. Lé vy flight search patterns of wandering albatrosses. Nature 381, 413–415 (1996).

[32] Viswanathan, G. M. et al. Optimizing the success of random searches. Nature 401, 911–914 (1999).

[33] Strogatz, S. H. Nonlinear Dynamics and Chaos: With Applications to Physics, Biology, Chemistry, and Engineering (CRC Press, 2018).

[34] Shaw, R. The Dripping Faucet as a Model Chaotic System (The Science Frontier Express Series) (Aerial Press, 1984).

[35] Gilpin, W., Bull, M. S. & Prakash, M. The multiscale physics of cilia and flagella. Nat Rev Phys 2, 74–88 (2020).

[36] Ullisch, I. A Closed-Form solution to the geometric goat problem. Math. Intelligencer 42, 12–16 (2020).

[37] Chupeau, M., Bénichou, O. & Voituriez, R. Cover times of random searches. Nat. Phys. 11, 844–847 (2015).

[38] Bhattacharjee, T. & Datta, S. S. Bacterial hopping and trapping in porous media. Nat. Commun. 10, 2075 (2019).

