## Supplemental Information for "Emergent Programmable Behavior and Chaos in Dynamically Driven Active Filaments"

June 5, 2022

**2 Supplementary Movie Captions** **6**

| Parameter name | Symbol | Value |
|---|---|---|
| No:of colloids | $N$ | 32 |
| Colloid radius | $a$ | 1 |
| Equilibrium bond length | $b_0$ | 2.1 |
| Axial spring stiffness | $k$ | 25 |
| Bending stiffness | $\kappa$ | 13.125 |
| Dipole strength | $d_0$ | 0.5 - 3 |

**Supplementary Table 1** Simulation parameters

### 1 Supplementary Text

#### 1.1 Simulation method and parameters

All simulations were implemented in Python with frequent and performance intensive functions implemented in Cython (an optimizing static compiler for Python [1]), to achieve significant performance gains. The mobility and propulsion tensors were calculated using the Pystokes library [2]. The governing equations of motion of the colloids were solved by integrating forward in time using backward-differentiation formulas implemented via the *Vode* function in the *odespy* python package [3]. All code used for generating the simulation data, analysis and figures presented in this paper can be found in the following Github repository: (https://github.com/deepakkrishnamurthy/PyFilaments). All simulations reported in the paper as well as analysis and data visualizations were done on a single desktop computer (MSI, CPU XYZ, RAM).

#### 1.2 Validation

##### 1.2.1 Euler-Bernoulli beam bending validation

We validated the bending module of the filament model using analytical results for Euler-Bernoulli beam bending [4]. We computed the final shape of a clamped filament given a transverse force acting at the free end and compared it with the analytical prediction and found good agreement (Fig. S1; Relative error: $< 2\%$ for 64 colloids for $\hat{\kappa} = 1250$ and

$F = 0.001$, $F = 0.01$ and $F = 0.05$).

##### 1.2.2 Filament flapping frequency under constant compressive load

To further validate the overall active filament model, we used our simulations to compute the flapping timescale (inverse of the flapping frequency) of a filament under a constant compressive tip follower-force and compared that to the predicted scaling law for this time-scale based on the underlying filament parameters which is given by [5]:

$$\tau_{flapping} = C\left(\frac{\mu L^4}{\kappa}\right)\mathcal{A}^{-4/3}, \tag{1}$$

where $\mu$ is the fluid viscosity, $L$ is the filament length, $\kappa$ is the bending rigidity and $\mathcal{A}$ is the dimensionless activity number and $C$ is an $\mathcal{O}(1)$ constant. As seen in Fig. S2 we find good agreement between the simulation prediction and the scaling law in Eq. (1).

#### 1.3 Fixed point analysis of the follower-force buckling map

Here we prove some results related to the fixed-points of the return-map of the filament orientation angle $\theta_{n+1} = f(\theta_n)$, which correspond to different types of limit cycles of the filament dynamics. We also derive the condition for the stability of these fixed-points/limit cycles.

##### 1.3.1 Fixed points corresponding to even iterate maps

First we prove that the fixed-points of $f(\theta) = -\theta$ correspond to period-2n limit cycles. We start with the observation that due to symmetry of the system about $\theta = 0$, the return map of filament orientation must be an odd-function of $\theta$:

$$f(-\theta) = -f(\theta). \tag{2}$$

Let $\theta^*$ be a fixed point of $f(\theta) = -\theta$, such that

$$f(\theta^*) = -\theta^*. \tag{3}$$

Now consider the second-iterate map $f^2(\theta) = f(f(\theta))$. At $\theta^*$, this is $f^2(\theta^*) = f(f(\theta^*)) = f(-\theta^*)$, from Eq.(3). But using Eq.(2) $f(-\theta^*) = -f(\theta^*)$, and from Eq.(3), $-f(\theta^*) = \theta^*$. Thus $\theta^*$ is a fixed-point of the second-iterate map.

In general, consider any even-iterate map $f^{2n}(\theta)$, where $n \in \mathcal{N}$. As shown above, if $\theta^*$ is a fixed point of $f(\theta) = -\theta$, then we have $f^{2n}(\theta^*) = (-1)^{2n-1} f(\theta^*) = \theta^*$, using the fact that for an odd function $f$, any function $f^m(\theta)$ is also odd. Thus if $\theta^*$ is a fixed point of $f(\theta) = -\theta$, then it is a fixed-point of any even-iterate map $f^{2n}(\theta)$.

##### 1.3.2 Stability of general fixed-points/limit cycles

The fixed points of $f(\theta) = \theta$ and $f(\theta) = -\theta$ correspond to limit cycles of the filament dynamics. The stability of these limit cycles can therefore be analyzed by looking at the stability of these fixed-points [6].

First we consider the fixed-points of $f(\theta) = \theta$, which correspond to 1-period dynamics. Let $\theta^*$ be a fixed point and $\eta_n$ be a perturbation about that fixed-point during cycle $n$, whose growth/decay is of interest over subsequent cycles. We have $\theta_n = \theta^* + \eta_n$ and $\theta_{n+1} = \theta^* + \eta_{n+1}$. We also have:

$$\theta^* + \eta_{n+1} = f(\theta^* + \eta_n), \tag{4}$$

which we expand in a Taylor series about $\theta^*$, to obtain:

$$\eta_{n+1} = f'(\theta^*)\eta_n + \mathcal{O}(\eta_n^2). \tag{5}$$

In general, we must have after $n$ cycles, $\eta_n = [f'(\theta^*)]^n \eta_0$, thus the dynamics is linearly stable if $|f'(\theta^*)| < 1$ and unstable if $|f'(\theta^*)| > 1$.

##### 1.3.3 Stability of even-period limit cycles

Now we consider the stability of the fixed-points of $f(\theta) = -\theta$, which correspond to even-period limit cycles of the dynamics. The growth/decay of a perturbation is governed by:

$$\theta^* + \eta_{2n} = f^{2n}(\theta^* + \eta). \tag{6}$$

Taylor expanding about $\theta^*$, we get:

$$\eta_{2n} = [f^{2n}(\theta^*)]'\eta + \mathcal{O}(\eta^2). \tag{7}$$

To linear order in $\eta$, the stability is given by $\lambda = [f^{2n}(\theta^*)]'$. We can simplify this using the chain-rule for differentiation:

$$\lambda = [f^{2n}(\theta^*)]' = f'(f^{2n-1}(\theta^*))f'(\theta^*) \tag{8}$$

$$\implies \lambda = f'(f^{2n-2}(f(\theta^*)))f'(\theta^*) \tag{9}$$

Using Eq.(3), Eq. (2) and the fact that $f^m(\theta)$ is odd if $f(\theta)$ is odd, we have:

$$\lambda = f'(-f^{2n-2}(\theta^*)) = f'(f^{2n-2}(\theta^*)), \tag{10}$$

since if $f$ is odd, then $f'$ must be even. Proceeding similarly, we finally obtain:

$$\lambda = f'(\theta^*)^2. \tag{11}$$

Thus an even-period cycle is linearly stable if $|f'(\theta^*)| < 1$ and unstable if $|f'(\theta^*)| > 1$, where $|f'(\theta^*)|$ is the magnitude of the slope at $\theta^*$ where $f(\theta^*) = -\theta^*$.

#### 2 Supplementary Movie Captions

**Movie S1** Flow around the filament during a cycle of compressional and extensional follower-force activity.

**Movie S2** Examples of periodic filament dynamics including 2, 6 and 10 periods.

**Movie S3** Examples of aperiodic filament behaviors for long times

**Movie S4** Periodic dynamics: Visualizing the evolution of nearby initial conditions in a reduced-order state-space.

**Movie S5** Chaotic dynamics: Visualizing the rapid divergence of nearby initial conditions as dynamics in a reduced-order state-space.

**Movie S6** Programming filament behaviors including (1) Homing and (2) Local vs Global search using frequency and amplitude modulated activity.

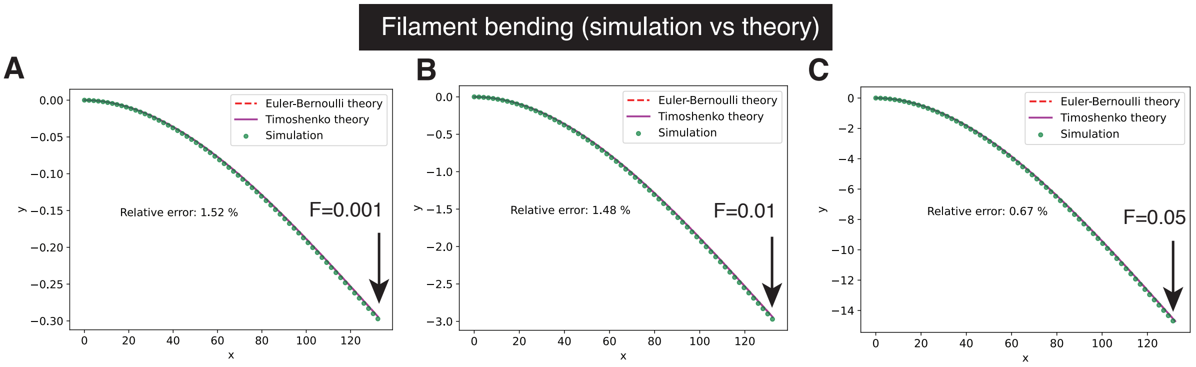

**Fig. S1 Validation: filament bending**. Validation of the filament model by comparing the computed shape due to a transverse tip force to analytical results from Euler-Bernoulli beam bending theory. (A) $F = 0.001$, (B) $F = 0.01$ adnd (C) $F = 0.05$ and $\kappa = 1250$, $N = 64$ colloids.

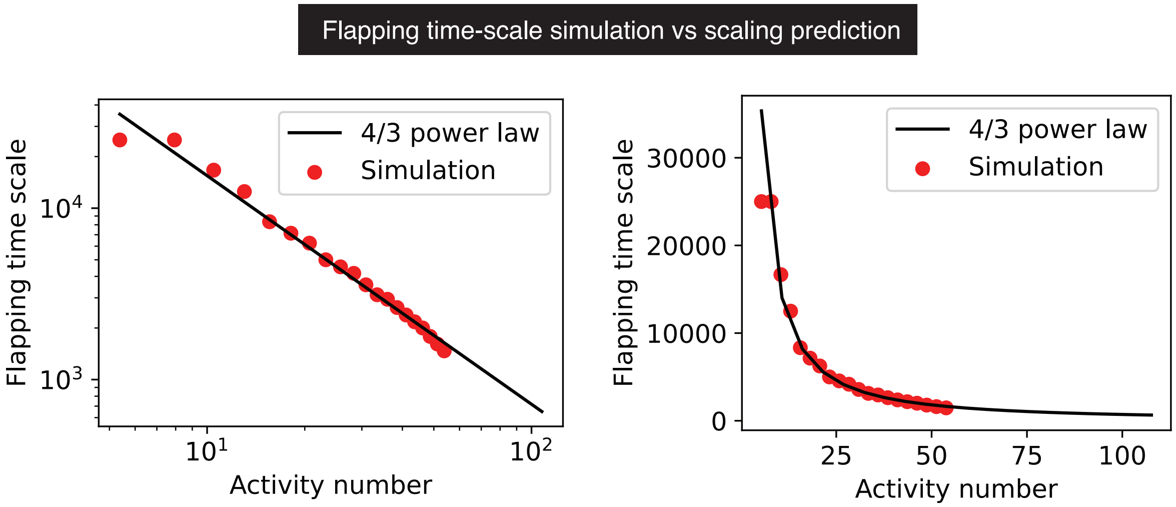

**Fig. S2 Validation: Filament flapping dynamics under constant compressive tip follower-force.** Comparing the flapping time-scale for a filament with constant, compressive tip follower force. Symbols our from our numerical experiments and the line shows the $4/3$ power-law from [7], shown in Eq. (1).

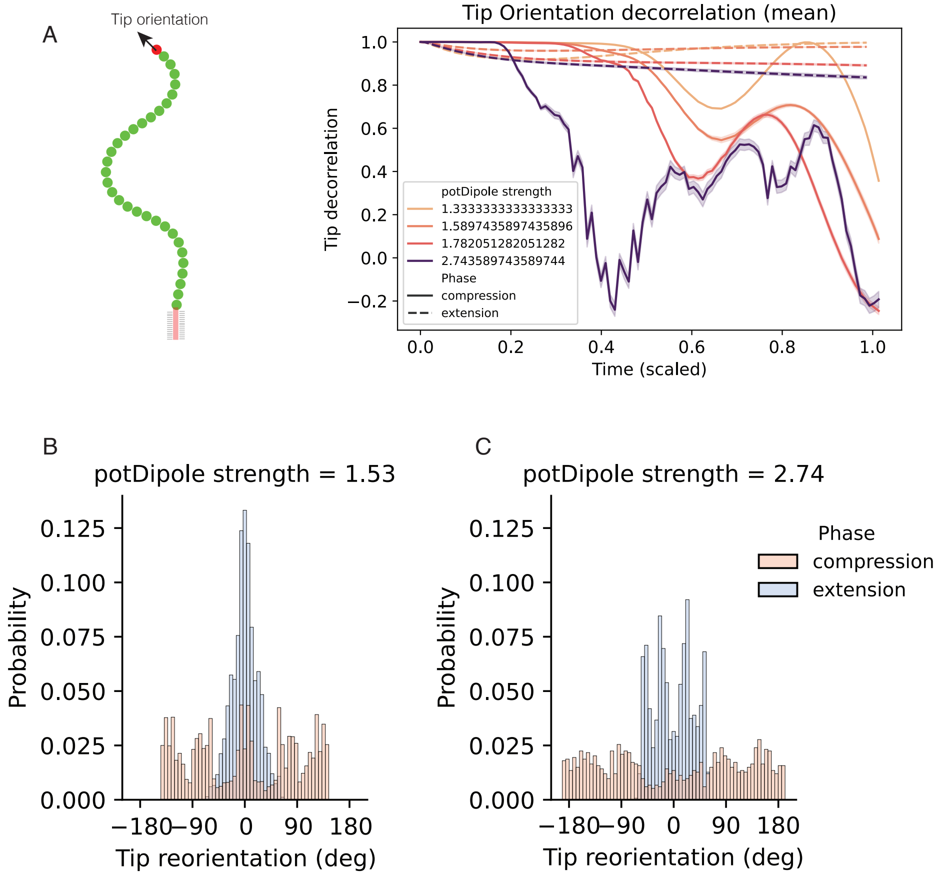

**Fig. S3 Compressive activity changes filament tip orientation while extension largely preserves it**. **(A)** Tip orientation decorrelation during compression (solid lines) and extension (dashed lines) for different values of activity strength. **(B)** and **(C)** Tip reorientation distribution for two different values of activity strength for compression and extension phases, shown in red and blue, respectively. The distributions show how compression scrambles tip orientation while extension largely preserved tip orientation.

### References

[1] Cython: C-Extensions for python. https://cython.org/. Accessed: 2022-5-20.

[2] Singh, R. & Adhikari, R. PyStokes: phoresis and stokesian hydrodynamics in python. *Journal of Open Source Software* **5**, 2318 (2020).

[3] Langtangen, H. P. odespy: Easy access in python to a large collection of ODE solvers.

[4] Gazzola, M., Dudte, L. H., McCormick, A. G. & Mahadevan, L. Forward and inverse problems in the mechanics of soft filaments. *Royal Society Open Science* **5** (2018).

[5] Chelakkot, R., Gopinath, A., Mahadevan, L. & Hagan, M. F. Flagellar dynamics of a connected chain of active, polar, brownian particles. *J. R. Soc. Interface* **11** (2014).

[6] Strogatz, S. H. *Nonlinear Dynamics and Chaos: With Applications to Physics, Biology, Chemistry, and Engineering* (CRC Press, 2018).

[7] De Canio, G., Lauga, E. & Goldstein, R. E. Spontaneous oscillations of elastic filaments induced by molecular motors. *J. R. Soc. Interface* **14** (2017).
